# The RNA binding protein LIN28A mediates chromatin dynamics during neuronal differentiation

**DOI:** 10.1101/2024.11.19.624261

**Authors:** Silvia Piscitelli, Emanuela Cascone, Chiara D’Ambrosio, Giuseppina Divisato, Laura De Lisio, Guido Leoni, Danilo Swann Matassa, Chiara Lanzuolo, Valentina Rosti, Maria Chiara Zizolfi, Monica Matuozzo, Emanuele di Patrizio Soldateschi, Paolo Maiuri, Andrea Scaloni, Fabiana Passaro, Silvia Parisi

## Abstract

The transition of embryonic stem cells (ESCs) from pluripotency to lineage commitment is regulated by multiple mechanisms, including chromatin dynamics and both transcriptional and post-transcriptional processes. Recent advances have highlighted that these mechanisms often interact, forming intricate multi-layered regulatory networks that require detailed characterization. In this study, we demonstrate that the RNA-binding protein LIN28A plays a pivotal role in neuronal differentiation by mediating RNA-dependent interactions with the Polycomb repressive complex 2 (PRC2). This interaction facilitates the eviction of PRC2 from chromatin, thereby activating a neuronal lineage-specific transcriptional program. Proteomic analyses revealed that the LIN28A interactome undergoes substantial remodeling during differentiation, corresponding to changes in LIN28A localization. In ESCs, LIN28A is predominantly nuclear and interacts with several components of the PRC2 complex in an RNA-dependent manner, assisting in chromatin dynamics. Our findings show that in the absence of LIN28A, PRC2 remains associated with chromatin, impairing the expression of genes critical for neuronal differentiation in ESCs. Chromatin immunoprecipitation sequencing (ChIP-seq) further confirmed that loss of LIN28A results in preferential PRC2 occupancy at the promoters of differentiation-associated genes. This study uncovers a novel role for LIN28A in epigenetic remodeling, which is essential for the proper differentiation of ESCs into the neuronal lineage.

## Introduction

Embryonic stem cells (ESCs), derived from the inner cell mass of the blastocyst, possess the unique abilities to self-renew and differentiate into all cell types comprising the three germ layers during development. These properties have traditionally been attributed to transcription factors such as octamer-binding transcription factor 4 (OCT4), sex-determining region Y-box 2 (SOX2), and the homeodomain-containing protein NANOG, as well as to chromatin-related factors (1). However, in recent years, post-transcriptional regulation by RNA binding proteins (RBPs) has emerged as an additional key mechanism controlling ESC pluripotency and early differentiation (2–4). Among RBPs, LIN28A stands out for its crucial role in regulating developmental timing, cellular proliferation, differentiation, stem cell pluripotency, and metabolism. Initially, LIN28A was identified as a negative regulator of the processing of precursor let-7 microRNAs (pre-let-7) into mature miRNAs (5–8). Subsequent studies revealed that LIN28A binds thousands of RNA transcripts, influencing their stability and translation efficiency (9–13).

LIN28A is highly expressed during embryogenesis and absent in most adult tissues (14). *In vitro*, LIN28A is present in the naïve state of ESCs, which resembles the pre-implantation embryo, and its levels increase in the primed state, which mimics the post-implantation epiblast. Loss of both LIN28A and its paralog LIN28B traps pluripotent stem cells (PSCs) in a more naïve state due to their function in translation of metabolic enzymes (15). Our group has demonstrated that during the transition of ESCs into epiblast-like cells (EpiLCs), LIN28A limits the translation of the architectural factor Hmga2 while enhancing that of the epigenetic regulator Dnmt3a, facilitating the proper exit of ESCs from the naïve state (10,11,16,17). LIN28A also plays an important role in the establishment of pluripotency. Remarkably, LIN28A is the only known RBP that, in combination with the Yamanaka factors (OCT4, SOX2, KLF4, and c-MYC), enhances the efficiency of reprogramming from somatic cells into induced pluripotent stem cells (iPSCs) (18–20). LIN28A achieves this by repressing the translation of mRNAs encoding proteins involved in oxidative phosphorylation and conferring to somatic cells the metabolic characteristics of PSCs (15).

We and others have observed that the effects of LIN28A depend not only on the RNAs it binds but also on its interactions with specific protein partners (9,11–13,21). Therefore, in this study, we have analyzed the interactome of LIN28A in undifferentiated ESCs and ESC-derived neural precursors. We have found a limited overlap of LIN28A interactors, suggesting different protein functions in undifferentiated and differentiated cells. Subunits of Polycomb repressive complex 2 (PRC2) in particular, are among ESC-specific interactors of LIN28A. PRC2 is a chromatin repressor complex responsible for trimethylating histone H3 at lysine 27 (H3K27me3) (22,23). The chromatin targeting of PRC2 and the regulation of its activity are very complex and still under investigation. Numerous studies have implicated RNA in the recruitment/eviction of PRCs from chromatin to regulate gene expression during development and differentiation (24–29). Here, we demonstrate that LIN28A binds PRC2 in an RNA-dependent manner, and this interaction facilitates the eviction of PRC2 from chromatin, enabling the expression of developmental genes during neural differentiation of ESCs.

## MATERIALS AND METHODS

### Cell culture, differentiation, treatments, and transfection

Mouse ESCs (E14Tg2a, BayGenomics) were grown on gelatine-coated plates in the following ESC medium: Glasgow minimum essential medium (Sigma-Aldrich) supplemented with 2 mM glutamine, 1 mM sodium pyruvate, 1]×]non-essential amino acids (all from Thermo Fisher Scientific), 0.1 mM β-mercaptoethanol (Sigma-Aldrich), 10% fetal bovine serum (Hyclone Laboratories), and 10^3^ U/ml leukaemia inhibitory factor (LIF, EMD Millipore).

The generation of EpiLCs was induced by plating ESCs on fibronectin coated plates in EpiLC medium: 1 vol of DMEM/F-12 combined with 1 vol of Neurobasal medium, supplemented with 0.5% N2 supplement, 1% B27 supplement, 1% KO serum replacement, 2 mM glutamine (all from Thermo Fisher Scientific), 20 ng/ml activin A (R&D Systems, Minneapolis, MN, USA), and 12 ng/ml basic fibroblast growth factor (Thermo Fisher Scientific). Within two days in these conditions, the cells express epiblast markers. We used the term “EpiLCs” to indicate cells at three days of differentiation from ESCs.

Differentiation toward neuroectodermal lineage was induced through SFEB formation by placing 1 × 10^5^ cells/ml in suspension in Petri dishes in the following differentiation medium: Glasgow minimum essential medium supplemented with 2 mM glutamine, 1 mM sodium pyruvate, 1× nonessential amino acids, 0.1 mM β-mercaptoethanol, and 10% KO serum replacement (Thermo Fisher). The term SFEBs indicates aggregates at 4 days of ESC differentiation unless noted otherwise. To obtain differentiation into neurons, 4 day-differentiated SFEBs were dissociated and plated on poly-D-lysine (Sigma-Aldrich)-coated plates in differentiation medium supplemented with 0.1 µM retinoic acid (Sigma-Aldrich). After 3-4 days, the appearance of neurite networks was already evident by morphological observation.

For the treatment with PRC2 inhibitor, ESCs were induced to differentiate into SFEBs in the presence or in the absence of 10 μM GSK126 (S7061, Selleckchem).

For the endodermal differentiation, 3 × 10^4^ cells/cm^2^ were plated on gelatin-coated plates in ESC medium and after 24 hours, the medium was replaced with RPMI (Gibco) supplemented with 1 μM Retinoic Acid, 3 μM CHIR-99021, 20ng/ml Activin, 10 ng/ml LIF, 2 mM glutamine and 2% of KSR. Cells were harvested at 4 days and 8 days of differentiation.

For cardiomyocyte differentiation we adopted the hanging drop method as previously described (30). Briefly, a suspension of 40.000 cells/ml in ESC medium without LIF was used to generate hanging drops of 20 μl/each to induce the formation of aggregates (embryoid bosied, EBs). After two days the EBs were collected in the same medium and grown in suspension for three days. Then, single EBs were placed in gelatin-coated well of 48-well plates and the counting of beating hearts was done at 12 days of differentiation.

For transfection ESCs were plated at 80.000 cells/cm^2^ on gelatin-coated plates. After sixteen hours ESCs were transfected with the plasmid pCAG-Lin28a-Flag plasmid expressing the FLAG-tagged form of LIN28A (10) or the empty vector, or with siRNAs (Thermo Fisher) using Lipofectamine 2000 (Invitrogen), according to the manufacturer’s protocol.

### Generation of Lin28 KO ESCs

To induce specific knock-out of Lin28a in ESCs we have used the SpCas9(BB)2A-Puro vector (PX459) described by Ran et colleagues (31). The two sgRNA guides were designed using CRISPOR tool. The targeting region of the two sgRNAs designed is reported in Supplementary Figure 1, the sequences are:

sgLin28#1 5’-CACCGCTGCGGCTCGTCTGCCGCT-3’

sgLin28#2 5’-CACCGTCTATGACCGCCCGCGCTG-3’

The sgRNA guides were cloned in the pSpCas9(BB)-2A-puro vector (Addgene) using BbsI (Thermofisher) and the plasmid was transfected in ESCs using Lipofectamine 2000. Correct targeting in the selected clones was assessed by PCR and sequencing.

### Cell fractionation and Western blot

For cell fraction experiment we adapted previous published protocols (32,33). Briefly, cells were resuspended in the following ice-cold buffer A: 10 mM HEPES pH 7.9, 10 mM KCl, 1.5 mM MgCl2, 0.34 M sucrose, 10% glycerol, 0.1% Triton X-100, 1 mM DTT, and 1x protease inhibitors (Sigma-Aldrich). Then, lysates were centrifuged, and the supernatant was stored as cytoplasmic fraction. The pellet was resuspended in 1X volume of buffer A and 10% of the sample was stored as total nuclear fraction. Then, the sample was diluted in buffer B (3 mM EDTA, 0.2 mM EGTA, 1 mM DTT, protease inhibitors, 0.5% NP-40), vortexed, and incubated for 60 minutes on ice. Sample was centrifuged and the supernatant was stored as nucleoplasmic fraction. Pellet from the previous step was then resuspended in 10X volume of B-SDS 1X lysis buffer (composed of 50 mM Tris-Cl [pH 7.5], 2 mM EDTA, 2% SDS) to yield the chromatin fraction. The fractions were sonicated for 5 minutes at high potency (30 seconds ON, 30 seconds OFF) until clarification and diluted in Laemmli 2X. For western blot analysis, the proteins were separated by SDS-PAGE under reducing conditions, transferred on PVDF membranes (Merk Millipore) and incubated with the following primary antibodies reported in **Supplementary Table S1**.

Whole cell extracts were obtained using 1X Ripa buffer (Thermo Scientific) supplemented with protease inhibitors (Sigma-Aldrich) and phosphatase inhibitors (Roche). After lysis, the samples were sonicated with Bioruptor Plus and centrifugated at 14000×g for 15 min to isolate total nuclear proteins. Total proteins were quantified using Bradford Protein Assay (Bradford Reagent, Bio-Rad). For western blot analysis, the proteins were separated by SDS-PAGE under reducing conditions, transferred on PVDF membranes (Merk Millipore) and incubated with the primary antibodies reported in **Supplementary Table S1**.

### Immunofluorescence, proximity ligation assay (PLA) and microscopy

Undifferentiated ESCs were grown on 8-well chambered coverslip (Ibidi) coated with gelatin s, fixed in 4% paraformaldehyde 15’ at room temperature (RT), washed three times in 1X PBS, permeabilized with 0.5% Triton X-100 15’ at RT and incubated with the primary antibodies (listed in **Supplementary Table S1**) 1 h, RT or 4°C, overnight. Then the cells were washed three times in 1X PBS, incubated with suitable secondary antibodies (Alexa Fluor, Thermo Fisher) for 1 h at RT, washed again and counterstained with DAPI (Calbiochem).

SFEBs were collected at the indicated differentiation day fixed in 4% paraformaldehyde and, after dehydratation with increasing percentages of ethanol, were embedded in paraffin and sectioned in 7-μm slices. After rehydration, permeabilization (0.2 % TX-100 for 5 minutes) and washing the slices were boiled in citrate buffer. Primary antibodies (listed in **Supplementary Table S1**) were incubated in 10 % FBS/1 % BSA/0.1 % Tween 20/1× PBS for overnight at 4°C followed by washing and secondary antibody hybridization. Nuclei were counterstained with Dapi (Calbiochem).

For PLA experiment, ESCs were plated at 5 x 10^5^ cells/well on gelatin-coated chambered coverslip (Ibidi). Then, cells were harvested after 24 hours, washed with 1X PBS and fixed with 4% paraformaldehyde for 10 min at room temperature. PLA were conducted according to the manufacturer’s protocol (NaveniFlex kit - Navinci). For antigen detection, the cells were incubated with the antibodies reported in **Supplementary Table S1**.

Images were captured with Leica Thunder Imaging System (Leica Microsystems) equipped with a LEICA DFC9000 GTC camera, lumencor fluorescence LED light source. 63X or 100X oil immersion objective were used to acquire Z-slice images. Small volume computational clearing was used to remove the background signal derived from out-of-focus blur.

For PLA quantitation, images were analyzed using ImageJ software (Fiji). After being imported via the Bio-Formats plugin, the images were segmented, and 2D structures larger than 0.01 μm² were subsequently automatically identified using the Analyze Particles command. Figures and statistical analyses related to the quantification of PLA images were conducted using R. The following packages were utilized: stringr, ggplot2, ggpubr, RColorBrewer, and ggpattern (**Supplementary Table S1)**. For statistical comparisons between experimental conditions, a non-parametric Wilcoxon test was employed to assess differences in means.

### RNA extraction, reverse transcription and real-time PCR

Total RNA from undifferentiated and differentiated ESCs, EpiLCs, EBs and SFEBs was extracted using Tri-Sure (Bioline), according to the manufacturer’s protocol. Approximately 1 µg of total RNA was used for reverse transcription using RevertAid Reverse Transcriptase (Thermo Scientific). Quantitative PCR was run on QuantStudio 7 Flex Real Time PCR System using Fast SYBR Green PCR Master Mix (Thermo Fisher Scientific). The expression of Gapdh mRNA was used as an internal control, using 2^-ΔΔCt method. Gene-specific primers used are reported in **Supplementary Table S1**.

### RNA immunoprecipitation

All immunoprecipitation experiments were performed using the following RIP protocol. Briefly, ESCs were transfected with pCAG-Lin28a-Flag plasmid or empty vector (mock) as control. After 48h of transfection undifferentiated cells were harvested and resuspended in polysome lysis buffer supplemented with protease inhibitors (Sigma-Aldrich) and phosphatase inhibitors (Roche). After centrifugation to remove cell debris, one milligram of total proteins was immunoprecipitated with anti-Flag affinity gel (Sigma-Aldrich) in the following NT2 buffer: 50 mM Tris–HCl pH 7.5, 150 mM NaCl, 1 mM MgCl2, 0.05% NP-40. To recover protein complexes for proteomic analysis and western blot, after washing, the anti-Flag affinity gel beads were resuspended in 2]×]Laemmli buffer and boiled for 5 min. To recover the RNA, after washing the beads were treated with proteinase K (10 mg/ml; Sigma-Aldrich) for 1 h at 55 °C to allow the release of bound RNA. RNA was purified using Tri-Sure (Bioline), and first-strand cDNA synthesis and qPCR were carried out as indicated above. The qPCR results were analyzed relating the Ct of each sample to the Ct of the input sample (DCt]=]Ct (input) – Ct (IP)) and then applying 2^DeltaCt comparative analysis.

For immunoprecipitation experiments with SFEBs, ESCs were transfected with pCAG-Lin28a-Flag plasmid and after 12 hours were induced to differentiate into SFEBs. The cells were collected at 4 days of differentiation and immunoprecipitation was performed as described above.

For the treatment with RNAse, the lysates were treated with RNAse A 50 µg/ml for 30 minutes at RT and then immunoprecipitated as described above.

### Proteomic analysis, protein identification and bioinformatics analysis

For proteomic analysis, immunoprecipitated samples from ESCs and SFEBs transfected with LIN28A-FLAG plasmid or empty vector (mock) as control were prepared as described in “RNA immunoprecipitation” section. Immunopurified proteins were analyzed by 12% T SDS-PAGE. After staining with colloidal Coomassie blue, whole gel lanes were cut into 15 slices, minced and washed with water. Corresponding proteins were separately in-gel reduced, S-alkylated with iodoacetamide and digested with trypsin, as previously reported (34). Individual protein digests were then analyzed with a nanoLC-ESI-Q-Orbitrap-MS/MS platform consisting of an UltiMate 3000 HPLC RSLC nano system (Thermo Fisher Scientific, USA) coupled to a Q-ExactivePlus mass spectrometer through a Nanoflex ion source (Thermo Fisher Scientific). Peptides were loaded on an Acclaim PepMapTM RSLC C18 column (150 mm × 75 μm ID, 2 μm particles, 100 Å pore size) (Thermo Fisher Scientific), and eluted with a gradient of solvent B (19.92/80/0.08 v/v/v water/acetonitrile/formic acid) in solvent A (99.9/0.1 v/v water/formic acid), at a flow rate of 300 nl/min. The gradient of solvent B started at 3%, increased to 40% over 40 min, raised to 80% over 5 min, remained at 80% for 4 min, and finally returned to 3% in 1 min, with a column equilibrating step of 30 min before the subsequent chromatographic run. The mass spectrometer operated in data-dependent mode using a full scan m/z range 375-1,500, a nominal resolution of 70,000, an automatic gain control target of 3,000,000, and a maximum target of 50 ms, followed by MS/MS scans of the 10 most abundant ions. MS/MS spectra were acquired in a scan m/z range 200-2000, using a normalized collision energy of 32%, an automatic gain control target of 100,000, a maximum ion target of 100 ms, and a resolution of 17,500. A dynamic exclusion value of 30 s was also used. Triplicate analysis of each sample was performed to increase the number of identified peptides/protein coverage.

MS and MS/MS raw data files per lane were merged for protein identification into Proteome Discoverer v. 2.4 software (Thermo Scientific), enabling the database search by Mascot algorithm v. 2.4.2 (Matrix Science, UK) with the following parameters: UniProtKB mouse protein database including the most common protein contaminants; carbamidomethylation at Cys as fixed modification; oxidation at Met, deamidation at Asn and Gln, and pyroglutamate formation at Gln as variable modifications. Peptide mass tolerance and fragment mass tolerance were set to ± 10 ppm and ± 0.05 Da, respectively. Proteolytic enzyme and maximum number of missed cleavages were set to trypsin and 2, respectively. Protein candidates assigned on the basis of at least two sequenced peptides and Mascot score ≥30 were considered confidently identified. Definitive peptide assignment was always associated with manual spectra visualization and verification. Results were filtered to 1% false discovery rate. A comparison with results from the corresponding control allowed to identify contaminant proteins that, nonetheless their abundance, were removed from the list of Lin28-interacting partners. A total of 5029 protein entries was identified. The mass spectrometry proteomics data have been deposited to the ProteomeXchange Consortium via the PRIDE (35) partner repository with a dataset identifier PXD049332. The list of LIN28A interactors was compared with those analyzed in different biological contexts (11,21), and BioGRID (36, IntAct (37) and STRING databases. Functional enrichment analysis including Gene Ontology was performed by using Metascape and FunRich 3.1.3 tools, setting *Mus musculus* and UniProtKB Rodents database, respectively. Venn diagrams were obtained by Bioinformatics & Evolutionary Genomics online tool. STRING online tool was used to visualize and integrate complex networks of proteomics data.

### Chromatin Immunoprecipitation sequencing (ChIP-seq)

Pellets of 3×10^6^ cross-linked ESCs or 2 days SFEBs were stored at -80°C until sonication. Each pellet was resuspended in 600 μL of cold Lysis Buffer: 50 mM Hepes-KOH, pH 7.5, 10 mM NaCl, 1 mM EDTA, 10% glycerol, 0.5% NP-40 and 0.25% Triton X-100. After 10 minutes on a rotator at 4°C, nuclei were collected by centrifugation for 5 minutes at 1350g at 4°C and resuspended in 130 µl of the following sonication buffer: 10 mM Tris-HCl pH 8.0, 2 mM EDTA, 0.1% SDS, 1X Protease Inhibitor Cocktail (Roche), 1 mM PMSF (Sigma-Aldrich). Cells were incubated on ice for 1 hour and total extracted chromatin was sonicated in a Covaris M220 focused-ultrasonicator using snap cap microTUBEs (Covaris) with cycles/burst 250, duration 18’. Fragmentation of chromatin to an average size of 200-500bp was checked on Agilent 2100 Bioanalyzer using High Sensitivity DNA Kit (Agilent). Samples were then diluted by adding 1 volume of equilibration buffer: 0.1% SDS, 10 mM Tris-HCl pH8, 233 mM NaCl, 1.66% Triton X-100, 0.166% DOC, 1 mM EDTA, 1X Protease Inhibitor Cocktail (Roche), 1 mM PMSF (Sigma-Aldrich2). To remove insoluble material samples were centrifuged at 14000g for 10 minutes at 4°C and supernatants were quantified using Nanodrop 1000 spectrophotometer. For each experimental condition 3% of total chromatin was stored at 4°C as input sample. For immunoprecipitation, 150 to 250 µg of chromatin were incubated overnight on a rotator at 4°C with Ezh2 antibody (Cell Signaling Technology). On the next day, protein G beads (Life Technology) were added to each sample and incubated on rotator for 3 hours at 4°C. Beads were then washed 10 minutes on rotator at 4°C twice with IP buffer, twice with high-salt IP buffer: 10 mM Tris-HCl pH8, 500 mM NaCl, 1.66% Triton X-100, 0.166% DOC, 0.1% SDS, 1 mM EDTA, 1X Protease Inhibitor Cocktail (Roche); 1 mM PMSF (Sigma-Aldrich); once with RIPA-LiCl buffer: (10 mM Tris-HCl pH 8.0, 1 mM EDTA, 250 mM LiCl, 0.5% DOC, 0.5% NP-40, 1X Protease Inhibitor Cocktail (Roche); 1 mM PMSF (Sigma-Aldrich); twice with 10 mM Tris-HCl pH 8.0. Crosslinking was reversed by incubating the beads and input at 65°C overnight with 100 µl of Elution buffer: 10 mM Tris-HCl pH 8, 0.5 mM EDTA, 300mM, 0.4% SDS. On the next day, all samples were diluted with 100 µl of 1X TE, treated with 2.5 U of RNAse cocktail (Ambion) at 37° for 120 minutes, followed by addition of 100 µg of Proteinase K (Invitrogen) at 55° for 120 minutes. DNA was then isolated using phenol/chloroform (Sigma-Aldrich) extraction and precipitation in cold ethanol. Pellets were suspended in nuclease-free water and quantified using Qubit 2.0 fluorometer with Qubit dsDNA HS Assay Kits (Invitrogen). The libraries were then prepared using the NEBNext Ultra II DNA Library Prep Kit for Illumina (NEB) and NEBNext Multiplex Oligos for Illumina (NEB). Library quantification and quality assay were done with TapeStation System. Libraries with distinct adapter indexes were normalized to a concentration of 2 nM, equimolarly pooled, and then loaded onto the Illumina NextSeq 2000 instrument. The sequencing was performed with a minimal target of 10 million reads for 100 bases in single-end mode on the Illumina NextSeq 2000 instrument at Ospedale Policlinico in Milan.

### NGS data processing

The high-throughput Ezh2 ChIP-seq data generated for this study are available in the NCBI GEO database with following accession number GSE282208. The sequences were demultiplexed with bcl2fastq. ChIP-seq sequencing lanes were merged with the help of GNU parallel software. Raw sequence reads were preliminary inspected with FASTQC software. Fastq files were aligned to the mm10 genome by using BWA-0.7.18 software (PMC2705234). PCR duplicates were marked and removed with Picard (v2.30.0) tool. Regions of EZH2 peaks enrichment were identified by MACS2 2.2.9.1 software (PMCID: PMC2592715) by selecting in each experimental sample only the peaks identified with a q-value cutoff < 0.05. Peaks detected in each biological replicate were intersected by using the intersectBed utility of bedtools 2.31 software (PMC2832824). For each condition only the peaks called in at least two out of the three experimental replicates were considered as real ChIPseq peaks. Metagene profiles reported in **Figure 5E** were obtained by using ngsplot 2.63 (PMCID: PMC4028082) with the following parameters “-G mm10 -R genebody -L 2000”. Peaks obtained in each condition were annotated by using the annotatePeak function included in ChIPseeker 1.41.3 package (PMID: 25765347) and specifying as source of annotation the org.Mm.eg.db included in BioConductor 3.20.0. Coverage tracks at specific genes of interest reported in **Figure 5H** were obtained with the ggcoverage 1.3.0 R package (PMCID: PMC10413535).

### Statistical analysis

FThe number of biological replicates of each experiment is indicated in the figure legends. The means of at least three independent experiments were used to calculate SD and to perform statistical analysis. All P values were calculated by Student’s t test using a two-tailed test and paired samples unless stated otherwise.

## Results

### Lin28a relocalizes during ESC differentiation

Lin28a was previously described by us and other groups as being expressed in the naïve (ESCs) with an increase when they undergo the transition into EpiLCs (**Figure 1A**) (10,38,39). However, if Lin28a expression is regulated also during the formation of the three germ layers remains unknown. To investigate this, we induced ESC differentiation into the three germ layers and confirmed our previous observations (10). We found that LIN28A expression peaks on day four of differentiation, when neural precursors begin to develop in serum-free embryoid bodies (SFEBs). LIN28A levels then rapidly decline as differentiation progresses into neurons (**Figure 1A**). In contrast, during differentiation into mesodermal **(Figure 1B)** and endodermal lineages **(Figure 1C)**, LIN28A expression gradually decreases and diverges from its interactor Ddx3x (**Figure 1A,B,C**). These observations suggest that LIN28A may utilize distinct functional partners depending on the differentiation context. Since, a recent study demonstrated that LIN28 exhibits dynamic temporal and spatial expression during murine embryo development (40), we explored the possibility that subcellular distribution of LIN28A also changes between undifferentiated and differentiated ESCs. Immunofluorescence analysis on undifferentiated cells showed that LIN28A is predominantly localized in the nucleus, with some detectable cytoplasmic signal. The nuclear localization displayed an apparent nucleolar pattern, consistent with prior reports (40). However, this pattern appeared to be reverted upon differentiation, with a prevalence of cytoplasmatic localization of LIN28A in SFEBs (**Figure 1D**). To validate these observations, we performed cell fractionation and confirmed that LIN28A protein levels in the nucleus and cytoplasm inversely correlated as ESCs were induced to differentiate (**Figure 1E**). Together, these findings suggested a possible role of LIN28A in neural lineage formation, highlighting its dynamic regulation during differentiation.

**Figure 1:**
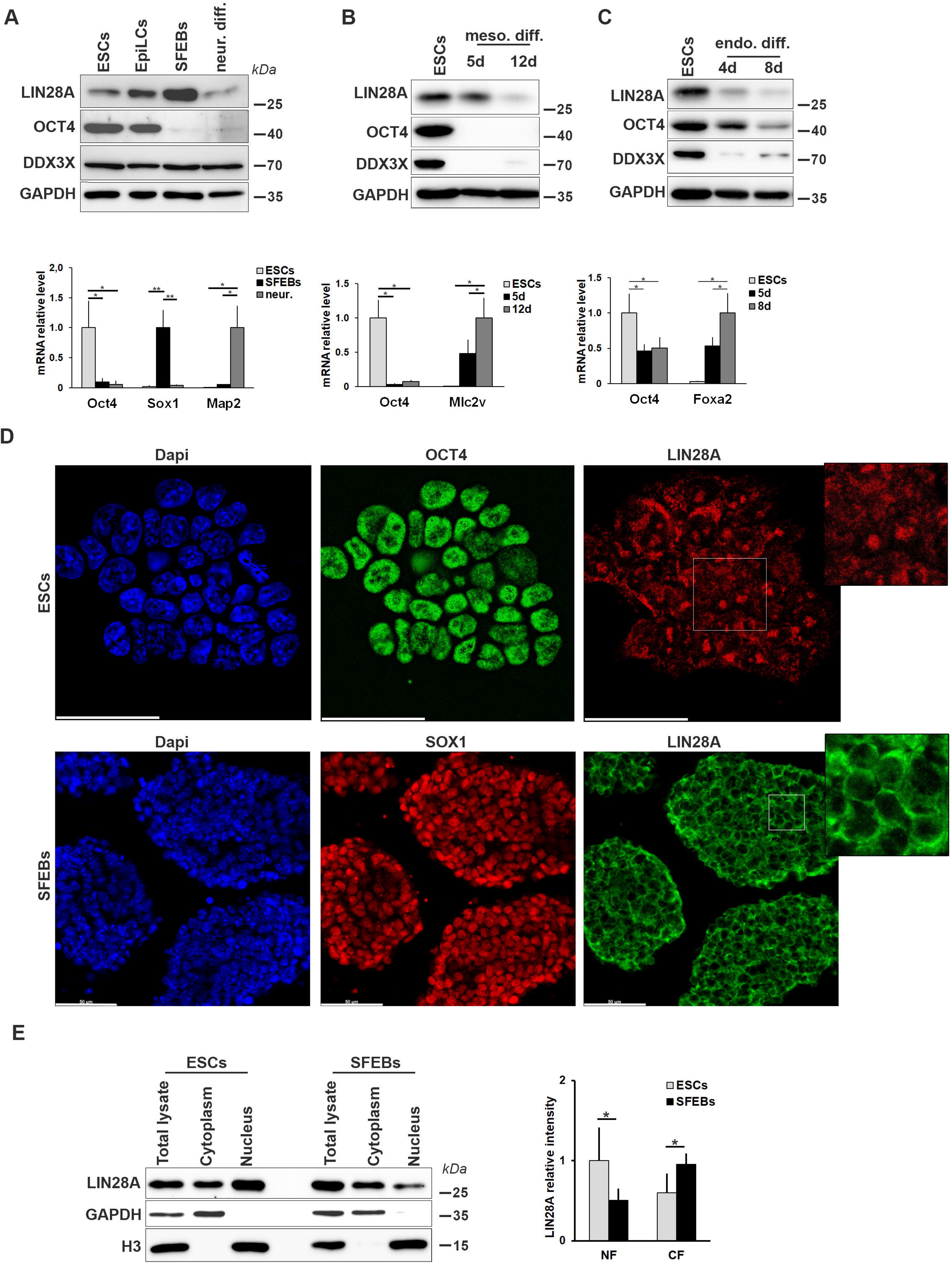
LIN28A expression profile and localization in undifferentiated and differentiated ESCs. A,B,C) Expression profile of LIN28A and its partner, DDX3X, in mouse ESCs induced to differentiate into EpiLCs, neural precursors (SFEBs) (A), mesodermal (B) and endodermal (C) lineages. The graphs under the blots represent the mRNA relative expression (using real-time RT-PCR) of the following specific markers: Oct4 for ESCs and EpilCs, Sox1 for neural precursors, Map2 for neurons, Mlc2v and Foxa2 for mesodermal and endodermal derivatives, respectively. Data are expresses as mean ± SD (n = 3), asterisks represent the statistical significance based on the Student’s t test (*p≤0.05, ** p≤0.005). C) Localization of endogenous LIN28A by immunofluorescence in undifferentiated ESCs and sectioned SFEBs. Oct4 and Sox1 were used as markers of undifferentiated ESCs and neural precursors, respectively. Scale bars: 50 μm. DAPI was used to counterstain the nuclei. Magnification of cropped areas of LIN28A staining are shown. D) Cell fractionation followed by western blot to evaluate the distribution of LIN28A in cell compartments of undifferentiated ESCs and SFEBs. The graph represents LIN28A levels relative to the quantity measured in the total lysate and normalized against GAPDH for the cytoplasmic fraction (CF) and against histone H3 for the nuclear fraction (NF). n = 5, *p≤0.05.

### LIN28A is required for neuronal differentiation

LIN28A has been reported to significantly improve the establishment of a pluripotent state during somatic cell reprogramming (20). However, its dynamic expression and localization changes during ESC differentiation suggesting a more specific role in neuronal lineage establishment. To investigate this possibility, we generated Lin28a knockout (KO) ESCs using CRISPR-Cas9 technology with two different single guide RNAs (sgRNAs) (**Supplementary Figure S1A**). Western blot and immunofluorescence analyses confirmed LIN28A protein depletion in both KO ESCs and SFEBs (**Supplementary Figure S1B,C and S2A**). These assays confirmed the efficacy of the sgRNAs and indicated that LIN28A loss is largely compatible with ESC viability in both undifferentiated and differentiated states (**Supplementary Figure S1B,C**). However, these initial analyses did not clarify whether LIN28A depletion affects ESC maintenance or differentiation potential. To evaluate the impact of LIN28A loss on ESC maintenance, we examined the ability of undifferentiated ESCs to form round alkaline phosphatase (AP)-positive colonies and observed no differences between WT and Lin28a KO cells (**Supplementary Figure S2A,B)**. Additionally, the percentage of undifferentiated ESCs expressing pluripotency markers was comparable between WT and KO cells (**Supplementary Figure S2C**). Finally, qPCR and western blot analyses over multiple passages revealed similar levels of pluripotency factors in WT and KO cells (**Supplementary Figure S2D, E**). Altogether, these results indicate that Lin28a depletion does not significantly impact the maintenance of undifferentiated ESCs. In contrast, when we induced differentiation into SFEBs, Lin28a KO cells displayed impaired neural differentiation. qPCR and western blot analyses revealed a failure to upregulate neural differentiation markers (Sox1 and Pax6) and a persistent expression of pluripotency markers (Oct4 and Nanog) (**Figure 2A,B)**. Consistently, immunostaining showed that less than half of Lin28a KO SFEB cells were Sox1-positive (∼30%) compared to WT cells (>80%). Conversely, a higher proportion (∼60%) of cells within KO SFEBs retained expression of Oct4 and Nanog (**Figure 2C**). Overall, these data indicated that Lin28a KO ESCs were not able to properly leave the undifferentiated state to develop into neural precursors.

**Figure 2:**
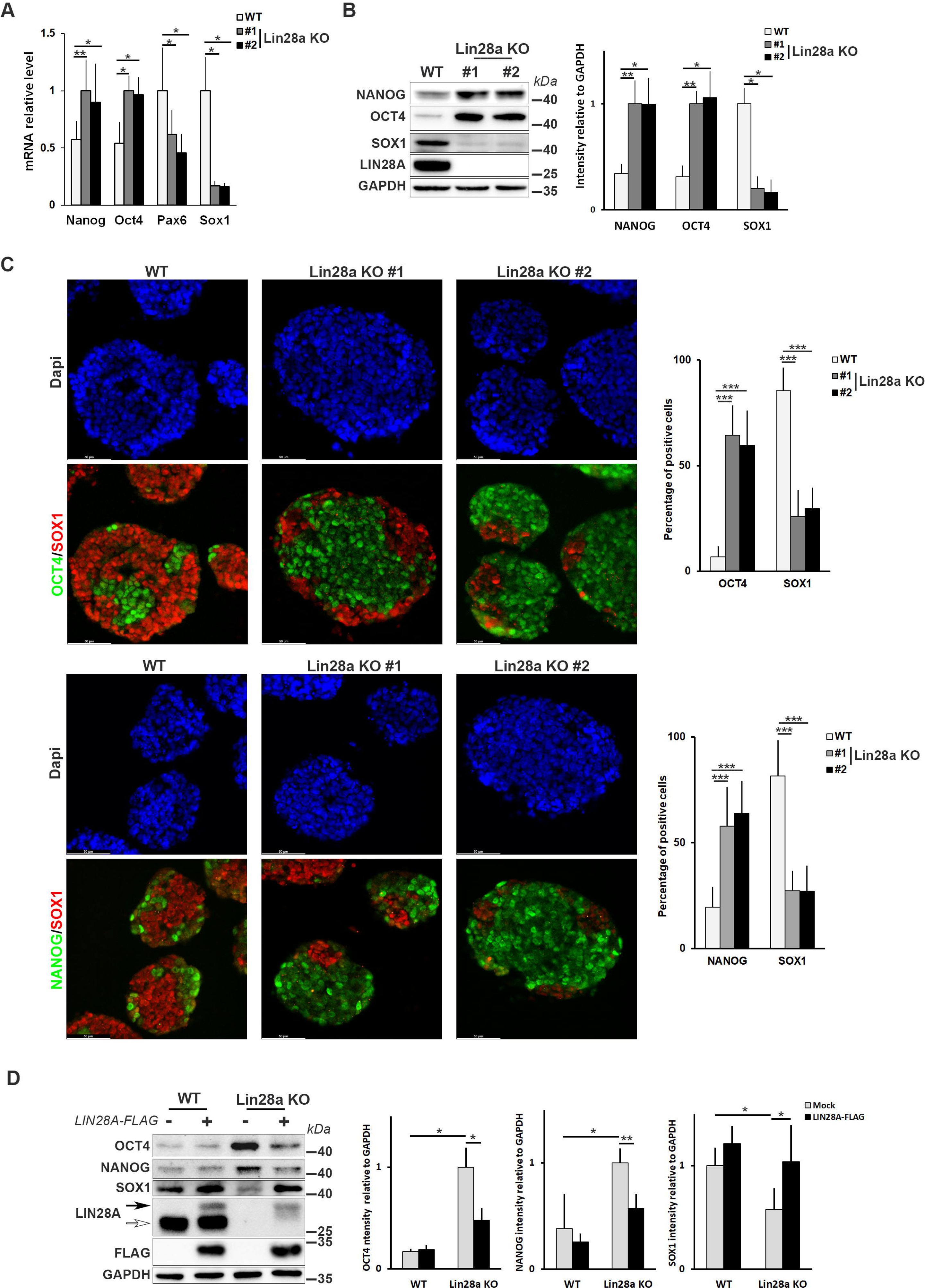
Lin28a KO ESCs are not able to properly differentiate. A) Expression levels of pluripotency (Oct4 and Nanog) and neural differentiation (Sox1 and Pax6) markers were evaluated by qPCR in WT and Lin28a KO SFEBs. n ≥3, *p≤0.05, ** p≤0.005. B) Western blot analysis of pluripotency and differentiation markers in WT and Lin28a KO SFEBs. The graph reports the relative intensity normalized for GAPDH. n = 4, *p≤0.05, ** p≤0.005. C) Immunofluorescence analysis of pluripotency (Oct4 and Nanog) and neural differentiation (Sox1) markers in sectioned SFEBs obtained from WT and Lin28a KO ESCs. DAPI was used to counterstain the nuclei. Scale bars: 50 μm. The graph represents the number of positive cells counting >400/cells and capturing 30 fields from three independent experiments. *** p≤0.0001. D) WT and Lin28a KO cells were transfected with the vector encoding a Flag-tagged form of LIN28A or the empty vector (Mock) and the differentiation into neural precursors was evaluated by western blot analysis using antibodies against pluripotency (Oct4 and Nanog) and neural (Sox1) Markers. The antibody against LIN28A shows the presence of both the endogenous protein (white arrow) and the FLAG-tagged form (black arrow). The graphs report the relative quantitation of three independent experiments. *p≤0.05; ** p≤0.0001.

To further confirm that this phenotype is specifically due to Lin28a, we transfected a Flag-tagged Lin28a construct in both WT and KO cells and induced differentiation. We found that the expression of exogenous Lin28a leads to downregulation of the pluripotency markers (Oct4 and Nanog) and upregulation of the neural differentiation marker Sox1 in the KO SFEBs (**Figure 2D and Supplementary Figure S3A).** These rescue experiments confirmed that the impaired differentiation phenotype is specifically attributable to Lin28a deficiency.

In order to evaluate whether differentiation defect in Lin28a KO cells is restricted to the neuroectodermal lineage or reflects a broader impairment, we analyzed the ability of KO cells to differentiate into mesodermal (i.e., beating cardiomyocytes) and endodermal lineages. Indeed, these analyses demonstrated that Lin28a KO ESCs differentiated into both lineages to the same extent of WT cells (**Supplementary Figure S3B,C**).

These findings, together with the prominent expression of Lin28a in neural precursors, demonstrated that this protein is required for the proper development of neural lineage, while being dispensable for mesendodermal lineage formation.

### Lin28a interactome changes during ESC differentiation

To elucidate the mechanistic role of LIN28A in the regulation of ESC fate, we analyzed the LIN28A interactome in both undifferentiated and differentiated cells. For this purpose, we expressed a Flag-tagged LIN28A in ESCs and induced differentiation into SFEBs. Then, we affinity-purified lysates from both ESCs and SFEBs using anti-Flag beads, followed by a gel-based proteomic approach to identify LIN28A protein interactors. Out of 5029 identified proteins, 816 were enriched in LIN28A-Flag ESCs compared to Mock ESCs (expressing the empty vector), and 364 protein entries were enriched in SFEBs relative to control immunoprecipitated (IP) samples (**Supplementary Table S2**). The specificity and sensitivity of this proteomic analysis were supported by a minimal presence of possible unspecific targets, according to the CRAPome database (41) (**Supplementary Table S3**).

To understand whether LIN28A interactions are context-dependent, we compared our dataset with previously reported LIN28A interactomes (11,21,36,37) (**Supplementary Table S3**). We found that among all Lin28A interactors identified so far, 377 (31.95%) were detected in at least one of the above-reported datasets, while 803 (68.05%) were unique to our dataset (**Figure 3A**). This underscores that LIN28A has a context-dependent interactome.

**Figure 3:**
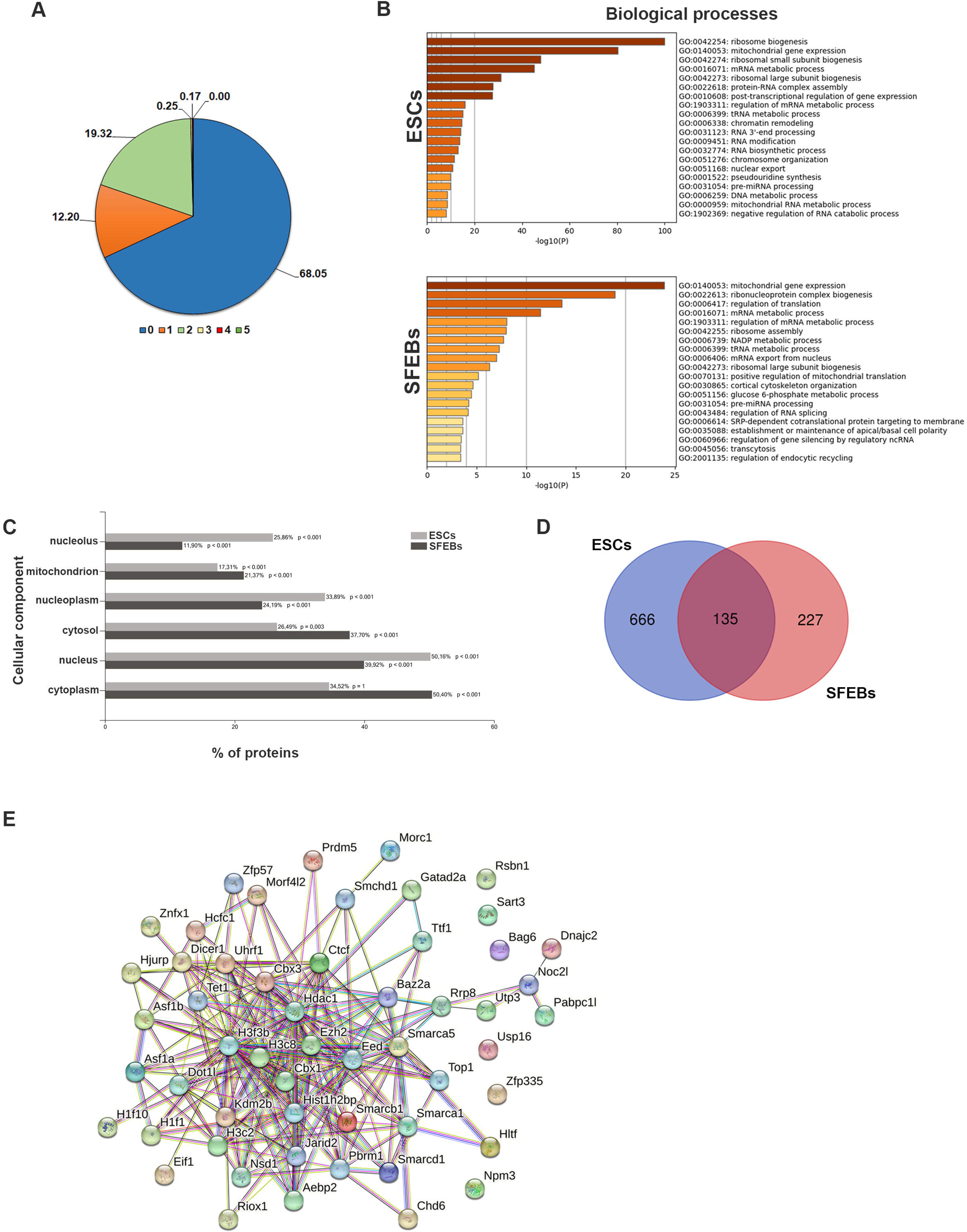
LIN28A changes interactors during ESC differentiation. A) Pie chart representing the percentage of putative LIN28A interactors identified in this study that were already described in recent studies. The highest percentage value (68.05) corresponded to LIN28A-binding proteins never reported before. We have reported also the percentage values of the interactors described once (12.20), twice (19.32), three times (0.25), four times (0.17) or five times (0.00) in the references and/or databases reported above. B) Metascape-based enrichment analysis for biological processes of the LIN28A interactors identified in ESCs or SFEBs. C) FunRich-based enrichment analysis for cellular components of the LIN28A interactors identified in ESCs or SFEBs. D) Venn diagram showing the high number of peculiar LIN28A interactors identified in ESCs and SFEBs, as well as the common proteins. E) STRING analysis detail of the proteins related to chromatin organization from LIN28A-interacting components uniquely identified in ESCs (medium-confidence, strength 0.49, False discovery rate 2.32 e-17).

To better understand the significant biological processes, cellular components, and the interconnections among the protein networks of these LIN28A-binding partners, we performed enrichment analyses. Gene ontology (GO) analysis of LIN28A-interacting partners in ESCs and SFEBs showed that, in agreement with previously published results, the most enriched biological processes were related to RNA processing, RNA metabolism and translation (**Figure 3B**) (7,9–11,13,15,42–44). On the other hand, GO terms in cellular component analysis showed that LIN28A interactors in ESCs are mainly localized in the nucleus, particularly in the nucleolus and nucleoplasm (**Figure 3B,C**). Conversely, in SFEBs, LIN28A interactors were primarily localized to the cytoplasm, cytosol, and mitochondria (**Figure 3B,C**), consistent with LIN28A’s subcellular distribution during differentiation (**Figure 1D,E**). Moreover, when comparing LIN28A interactomes across the two states, we found that 666 were specific for ESCs and 227 were unique to SFEBs, while only 135 were shared (**Figure 3D**). Interestingly, this indicates that LIN28A forms distinct molecular complexes in undifferentiated versus differentiated states, most likely exerting different functions therein.

Further GO analysis of ESC-specific interactors linked them to chromatin organization, RNA processing, metabolism and ribonucleoprotein complex biogenesis (**Supplementary Figure S4A**), aligning with LIN28A localization in the nucleus and in the nucleolus (**Figure 1D**). On the other hand, GO enriched terms in SFEBs were associated with protein transport, localization and cellular metabolic processes (**Supplementary Figure S4A**). Interestingly, GO analysis for molecular function of ESC-specific LIN28A interactors were enriched for DNA/RNA binding, while SFEB-specific interactors were associated with protein and small molecule binding (**Supplementary figure S4B**). To explore these interactions further, we constructed a protein-protein interaction (PPI) network using the STRING database. The shared interactors formed a small cluster related to mitochondrial gene expression and translation, suggesting an unknown LIN28A role in mitochondrial translation (**Supplementary Figure S4C**). The ESC-specific interactome formed a large, complex network with subclusters related to chromatin organization and remodeling (**Supplementary Figure S4D**), with 88 and 28 entries, respectively (**Figure 3E and Supplementary Figure S4E**). This observation suggests that LIN28A may exert its function in the determination of ESC fate through chromatin regulation, a role that remains only partially explored, and that represents the focus of our subsequent analyses.

### LIN28A influences epigenetic regulation

The unique interactions of LIN28A found in undifferentiated cells might explain why Lin28a KO cells fail to properly differentiate into neural precursors. Among ESC-specific LIN28A-binding partners, we found epigenetic regulators involved in chromatin remodeling including components of Polycomb Repressive Complex 2 (PRC2), the histone-modifying complex whose activity is associated with transcriptional silencing during ESC fate specification (45). Specifically, we identified the PRC2 core components EED (embryonic ectoderm development) and EZH2 (enhancer of zeste 2), as well as the accessory proteins AEBP2 (adipocyte enhancer-binding protein 2) and JARID2 (Jumonji and AT-rich interaction domain 2). While another PRC2 subunit, SUZ12 (suppressor of zeste 12), was initially filtered out due to a cutoff threshold, it was later validated. To confirm that the results from our IP-mass spec experiments could be validated in alternative assays, we performed again co-IP assays on lysates of ESCs ectopically expressing the FLAG-tagged LIN28A, but this time we analyzed the immunoprecipitates directly by western blot for the presence of PRC2 subunits. In agreement with proteomic results, the PRC2 component analyzed Ezh2 and Suz12, and Jarid2, were co-immunopurified with the FLAG-LIN28A **(Figure 4A**). Additionally, the proximity ligation assay (PLA) revealed the *in situ* interactions of LIN28A with the same PRC2 components (**Figure 4B**). Indeed, prominently higher levels of PLA-positive spots were observed between LIN28A and PRC proteins compared to P-Smad1/5, which was used as a negative control since it was not detected through proteomic experiments (**Figure 4B**).

**Figure 4:**
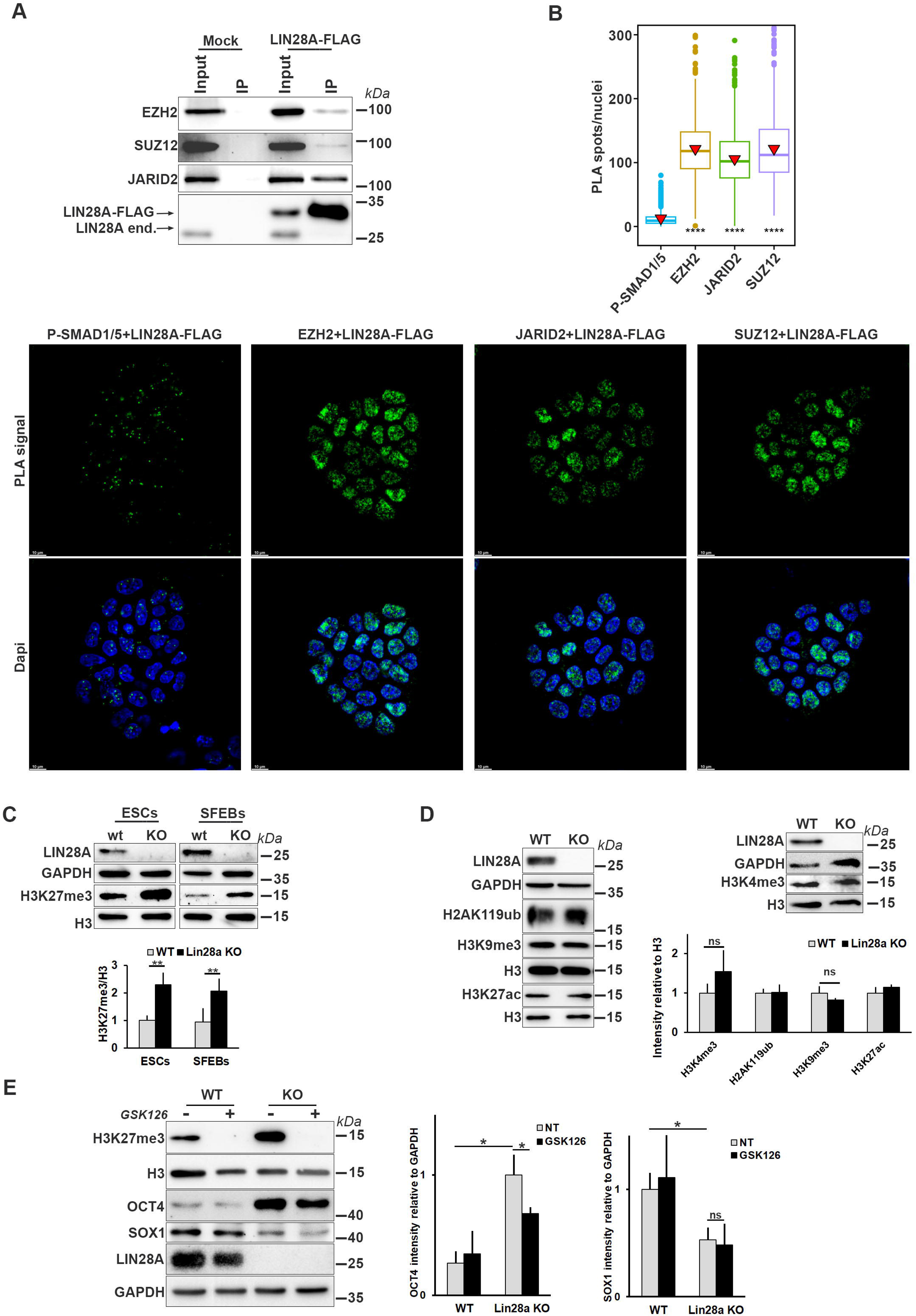
LIN28A binds to the PRC2 complex and influences its epigenetic activity. A) Western blot analysis of LIN28A immunoprecipitates showing the interaction of LIN28A with PRC2 subunits. Immunoprecipitation was performed using anti-FLAG beads in ESCs transfected with LIN28A-FLAG expressing vector or empty (Mock) vector. B) Proximity ligation assay (PLA) to assess the interaction of LIN28A with PRC2. ESCs expressing LIN28A-FLAG were analyzed by PLA coupling the anti-FLAG antibody with the following antibodies: anti-P-SMAD 1/5 (as negative control), anti-EZH2, anti-SUZ12 and anti-JARID2. In the corresponding images, PLA positive spots are visible in green. The quantification of positive spots/cell from three independent experiments is shown in the graph. ***p≤0.0001. C) Western blot analysis and relative quantification of H3K27me3 histone mark in WT and Lin28a KO cells in both undifferentiated and differentiated state. Histone H3 and GAPDH were used as loading control. The use of an antibody against LIN28A confirms the absence of protein in Lin28a KO cells. The graph represents the quantification of six independent experiments. **p < 0.005, ns: not significant. D) Western blot analysis and relative quantification of different histone marks in WT and Lin28a KO cells. Histone H3 and GAPDH were used as loading control. n=3, ns: not significant. E) Western blot analysis and relative quantification of the levels of pluripotency (OCT4) and differentiation (SOX1) markers in WT and Lin28a KO SFEBs treated with the PRC2 inhibitor, GSK126 during differentiation. GAPDH and histone H3 were used as loading control. n=3 biological replicates, *p < 0.05.

Given that the main function of PRC2 is to deposit trimethylation on lysine 27 of histone H3 (H3K27me3) (45), and with the aim of understanding whether LIN28A could influence the activity of the PRC2 complex, we evaluated this epigenetic mark in WT and Lin28a KO cells. We found an accumulation of H3K27me3 in undifferentiated KO ESCs compared to WT, which persisted even in differentiated cells (**Figure 4C**). Despite comparable expression levels of EZH2 in WT and KO cells **(Supplementary Figure S5A,B)**; we hypothesized that the observed increase of H3K27me3 deposition in Lin28a KO cells might be due to changes in protein activity or distribution on chromatin. ESCs are characterized by high presence of bivalent promoters decorated by both activating H3K4me3 mark and repressive H3K27me3 mark (46,47). To investigate whether the H3K4me3 mark parallels the accumulation of the H3K27me3 counterpart in Lin28a KO ESCs, we performed western blot analysis, but no significant differences in H3K4me3 levels were observed between WT and KO cells (**Figure 4D**). Next, given that the trimethylation of H3K27 occurs following the ubiquitination of H2AK119, a process catalyzed by Polycomb Repressive Complex 1 (PRC1) (45), we analyzed whether the increase of H3K27me3 observed in Lin28 KO cells was in turn associated with hyperactivity of PRC1. We observed no changes in H2AK119ub mark between Lin28a KO and WT ESCs (**Figure 4D**), further suggesting that the effects of Lin28a depletion specifically affects PRC2 activity. To determine whether Lin28a KO ESCs are also characterized by the alteration of other epigenetic marks, we analyzed the levels of other histone modifications. As shown in **Figure 4D**, none of the other epigenetic marks analyzed were altered in the absence of LIN28A. All together, these results indicated that the absence of Lin28a alters the level of PRC2 specific mark (H3K27me3), rather than causing a general change in the epigenetic state. This points to PRC2 as a likely culprit in the inability of Lin28a KO cells to properly differentiate. PRC2 is well-known for its role in establishing cell type-specific gene expression programs that enable differentiation. Therefore, since Lin28a KO cells are unable to properly differentiate, and show high H3K27me3 levels, we asked whether inhibition of PRC2 enzymatic activity might rescue the Lin28a KO cell phenotype. To explore this hypothesis, we treated WT and KO differentiating cells with GSK126, a highly selective inhibitor of EZH2 methyltransferase. Analysis of pluripotency and differentiation markers revealed that the reduction in H3K27me3 deposition induced by GSK126 only partially rescued the phenotype of Lin28a KO cells (**Figure 4E**). In fact, downregulation of the pluripotency marker Oct4 was at least in part restored, whereas the activation of the neuronal marker Sox1 remained impaired. These results indicate that the differentiation block observed in Lin28a KO ESCs cannot be fully attributed to the enzymatic activity of PRC2.

### LIN28A can modulate PRC2 binding at chromatin during differentiation

Since changes in PRC2 enzymatic activity alone cannot account for the inability of Lin28a KO ESCs to differentiate into neural precursors, we hypothesized that LIN28A can modulate the dynamics of PRC2 complex during differentiation. To test this hypothesis, we evaluated the amount of PRC2 associated with the chromatin in WT and Lin28a KO ESCs. As expected, we found that not all of PRC2 is associated with the chromatin fraction in WT ESCs (**Figure 5A**), consistent to a dynamic balance between PRC2 chromatin occupancy and its association to RNAs (25,26,48,49). However, we found that the fraction of Ezh2 associated with chromatin is higher in Lin28a KO cells compared to WT counterparts, particularly in differentiating ESCs (**Figure 5A,B**). This finding suggests that LIN28A, by binding RNA, could compete with Ezh2 and facilitate its eviction from chromatin during differentiation. PRC2 has been previously reported to be recruited to and displaced from Polycomb target genes during ESC differentiation (50). Therefore, we hypothesized that LIN28A could play a role in this dynamic during ESC differentiation. To test this idea, we performed chromatin immunoprecipitation for Ezh2 followed by deep sequencing (ChIP-seq) in undifferentiated and differentiating cells (2 days of SFEB differentiation) to evaluate the association and distribution of PRC2 complex to specific genomic loci in the presence or absence of LIN28A. We identified 4859 ChIP-seq peaks in WT ESCs and 2516 in Lin28a KO ESCs (**Figure 5C**). In differentiating cells, PRC2 occupancy was more extensive in Lin28a KO cells compared to WT (6338 versus 4495 EZH2 peaks) (**Figure 5C**). The majority (≥75%) of Ezh2-binding peaks were located at gene promoters (within 1000 bp from transcriptional start site, TSS and transcriptional end site, TES), with EZH2 signal enriched at TSSs under all conditions analyzed (**Figure 5D**). Meta-gene profiles of the detected EZH2 peaks highlighted an enrichment of the EZH2 signal in KO SFEBs compared to WT counterpart, whereas undifferentiated ESC KO and WT cells displayed quantitatively similar profiles (**Figure 5E**). Next, we analyzed the landscape of EZH2 target genes in WT and Lin28KO cells by associating to each detected peak the nearest gene with a TSS within +/- 1000 bp of distance (**Supplementary Table S4**). In undifferentiated cells almost all the genes that mapped in KO cells were common with those in WT cells. Conversely, in differentiating cells we found that about 30% of the mapped genes (1441 out of 4718) were bound only in KO cells (**Figure 5F**). This observation supported the evidence of a differential Ezh2 genome occupancy in differentiating WT and KO cells. Of note, GO enrichment analysis of the 1441 EZH2 targets found exclusively in differentiating Lin28a KO cells revealed that most of these genes are involved in neuronal system development, neuron differentiation and synapsis transmission (**Figure 5G**). Furthermore, EZH2 occupancy is enriched specifically in Lin28a KO differentiating cells at promoters of genes involved in neuronal differentiation (Pax6, Sall2, Tead3 and Cbx2) (**Figure 5H**). The persistent PRC2 occupancy at these promoters was accompanied by their transcriptional repression in KO cells, compared to WT (**Figure 5I**), consistent with the impairment of neuronal differentiation described above.

**Figure 5:**
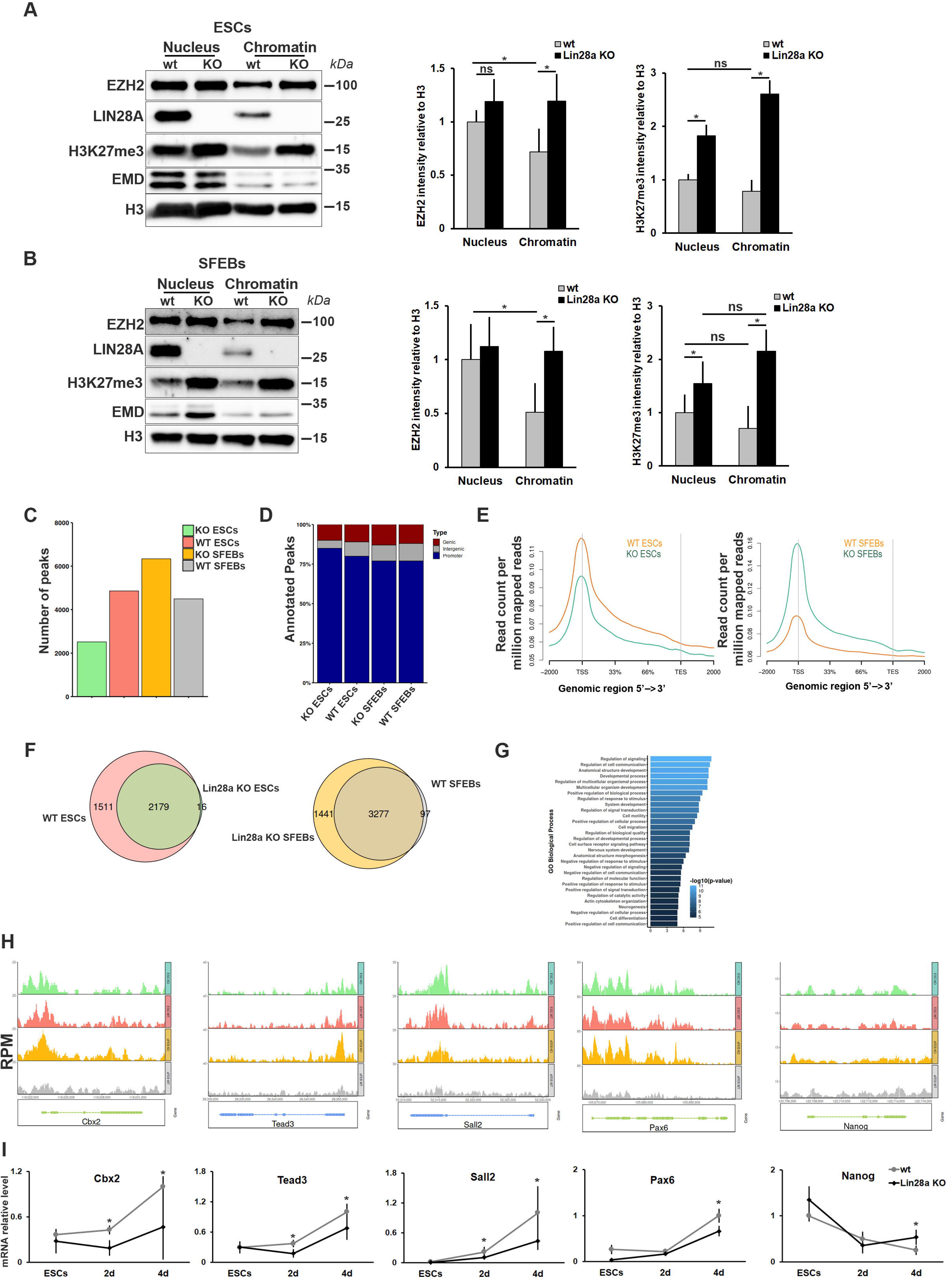
LIN28A influences PRC2 association with chromatin. A) Western blot analysis of EZH2 in total nuclear fraction and chromatin fraction of WT and Lin28a KO ESCs. B) Western blot analysis of EZH2 in total nuclear and chromatin fraction of WT and Lin28a KO at 2 days of SFEB differentiation. In both panels (A and B) GAPDH and H3 were used as loading control. Emerin was used as nuclear marker. The graphs represent the levels of EZH2 and H3K27me3 normalized to H3 signal in the relative fractions. n≥4, *p < 0.05, ns: not significant. C) Bar-plot reporting the number of peaks detected in each tested cell line by analyzing the ChIP-seq NGS experiments with MACS2 software. For each condition, only the peaks detected in at least 2 out of the 3 biological replicates analyzed were reported. The analysis detected 2516 peaks in Lin28a KO ESCs, 4859 in WT ESCs, 6338 peaks in Lin28a KO SFEBs, and 4495 peaks in WT SFEBs. D) Bar-plot reporting the distribution of peaks at promoters, genic or intergenic regions. (Promoter: peaks within +/- 1000 bp from the TSS; genic: peaks outside the promoter region but within a gene, intergenic outside annotated genes). E) Metagene profiles of EZH2 peaks. Intensity plot (reads per million) showing EZH2 ChIP-seq intensity in WT and Lin28a KO differentiated or undifferentiated cells at gene bodies (within 2000 bp). TSS: transcription start site; TES: transcription end site. F) Venn diagrams showing the overlap of the genes mapped on detected peaks (+/-1 1000 bp) in WT and Lin28a KO ESCs or SFEBs (complete list in **Supplementary Table S4**). G) Diagram reporting the GO for the 1441 genes enriched for EZH2 peaks specifically in Lin28a KO SFEBs (Benjamini Hochberg corrected p-value <=0.05). The top 30 significantly enriched biological processes and the relative p-value are reported H) EZH2 ChIP-seq tracks at the promoters of differentiation genes. Plots highlight the fingerprint of binding of EZH2 at the promoter of selected genes (Cbx2, Tead3, Sall2 and Pax6) and the pluripotency gene Nanog (as control). Intensity of the peaks is reported as reads per million. I) Expression profile of some differentiation genes on which EZH2 occupancy was enriched at promoters specifically in Lin28a KO SFEBs. The graphs report the expression levels at three time-points: undifferentiated ESCs (t0) 2 and 4 days after the induction of differentiation through SFEBs. Nanog was used as control. n=4, *p≤0.05.

These results demonstrated that, in Lin28a KO cells, EZH2 remains more associated with chromatin at specific developmental gene loci involved in neuronal development and differentiation. These results, together with the observation that the inhibition of PRC2 enzymatic activity (**Figure 4E**) cannot completely rescue the differentiation of Lin28a KO cells, support the idea that LIN28A is required to properly dissociate PRC2 from developmental gene loci during differentiation rather than regulating its enzymatic activity.

### LIN28A interaction with PRC2 is RNA-dependent

The association of PRC2 with mRNAs and/or non-coding RNAs (ncRNAs) is critical to regulate its recruitment to specific gene loci (26,51,52). To investigate whether LIN28A indeed regulates the recruitment of PRC2 to specific gene loci through its RNA-binding capacity, we treated lysates of cells expressing LIN28A-Flag with RNase and analyzed the interaction between LIN28A and PRC2 subunits by co-immunoprecipitation. We found that RNase treatment disrupted LIN28A’s ability to efficiently capture PRC2 components (EZH2, SUZ12 and JARID2), thus indicating that the physical association between LIN28A and PRC2 is RNA-dependent **(Figure 6A).**

**Figure 6:**
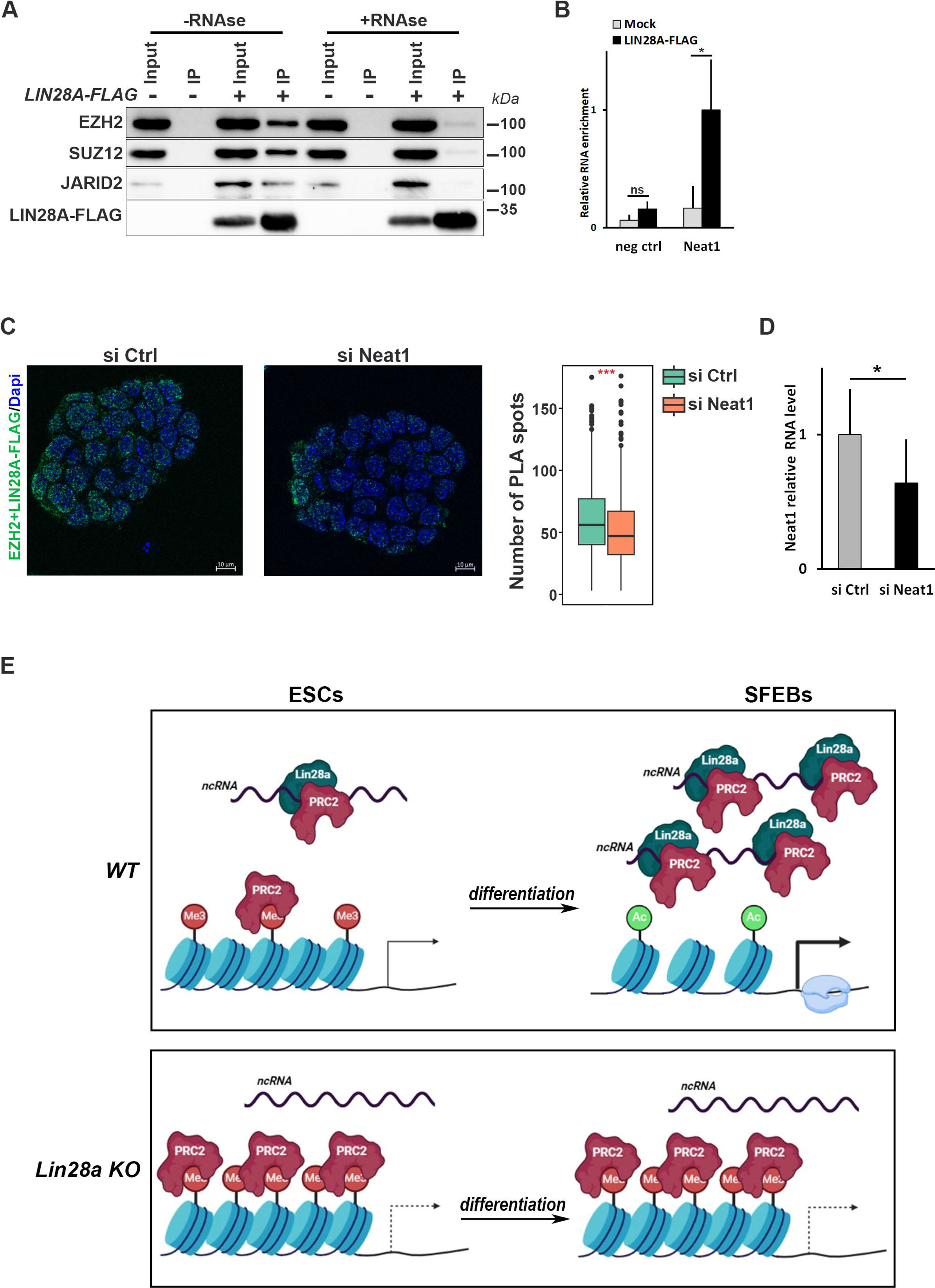
LIN28A interaction with PRC2 is RNA-dependent. A) Western blot analysis of LIN28A immunoprecipitates with or without RNAse treatment. Immunoprecipitation was performed using anti-FLAG beads in ESCs transfected with LIN28A-Flag expressing vector or empty vector. B) RNA immunoprecipitation followed by qPCR analysis to evaluate the binding of ncRNA Neat1 to LIN28A. The 3’UTR of Lin28b was used as negative control. n=4, *p < 0.05. C) Proximity ligation assay (PLA) to assess the interaction of LIN28A with EZH2 upon silencing of the ncRNA Neat1. ESCs expressing LIN28A-FLAG were analyzed by PLA coupling the anti-FLAG with anti-EZH2 antibodies. The quantification of positive PLA spots/nucleus and the mean area of spots from three independent experiments is shown in the graph. **p≤0.0001, ***p≤0.0001. D) qPCR analysis of Neat1 levels upon silencing with a specific siRNA for the experiments reported in the panel (C). n=3, *p≤0.05. E) Proposed model for LIN28A involvement in epigenetic regulation. In WT ESCs LIN28A counteracts PRC2 recruitment to chromatin allowing the activation of developmental genes upon differentiation. In Lin28a KO ESCs the PRC2 eviction from the chromatin is impaired because of the absence of LIN28A, and thus PRC2 occupancy blocks the activation of developmental genes and, in turn, cell differentiation.

Recent studies have identified the long non-coding RNA nuclear paraspeckle assembly transcript 1 (Neat1) as a key regulator of PRC2 activity. Neat1 is essential for PRC2 eviction from chromatin and activation of lineage-specific genes, potentially through interactions with yet unidentified RBPs (26). To understand whether LIN28A is one of these RBPs, we analyzed its binding to Neat1 through RNA immunoprecipitation and found that LIN28A efficiently binds to Neat1 RNA in ESCs **(Figure 6B)**. We next investigated whether the interaction between LIN28A and PRC2 could be mediated by Neat1. Using the PLA approach, we analyzed the spatial proximity of LIN28A and EZH2 upon Neat1 silencing in ESCs. As shown in **Figure 6C**, quantification of PLA-positive spots per nucleus revealed a significant reduction in LIN28A-EZH2 proximity even if Neat1 is partially silenced (**Figure 6D**). While the involvement of other molecules (RNAs and/or proteins) cannot be ruled out, these results strongly suggest that Neat1 serves as a potential scaffold facilitating the formation of the PRC2/LIN28A complex.

## Discussion

Beyond its role in miRNA biogenesis, the RNA-binding protein LIN28A has already been described as a key regulator of translation in ESCs (6,8,9,13). During the transition from ESCs into EpiLCs, LIN28A regulates the translation of the mRNAs of two important epigenetic players: the DNA methyltransferase DNMT3A and the architectural factor HMGA2 (10,11). Here, we demonstrate a novel role for LIN28A as epigenetic regulator of neural differentiation of ESCs.

Generation of Lin28a KO ESCs revealed that LIN28A, while dispensable for ESC self-renewal, is essential for proper differentiation into neuronal precursors, particularly in the early stages of this process. Protein localization and interactomics experiments in both undifferentiated and differentiated ESCs indicated that LIN28A undergoes changes in cellular localization and molecular interactions following differentiation. Our proteomic data align with LIN28A’s established roles in ribosome biogenesis, rRNA processing, and nucleolar remodelling and translation (9–13,40, 53). We have highlighted that only few interactor proteins are shared between undifferentiated and differentiated ESCs. Notably, over 13% f the interactors specific to undifferentiated ESCs (88 proteins) are associated with chromatin organization.

Zeng and coworkers previously proposed a role of LIN28A in epigenetic regulation, showing its recruitment of the 5-methylcytosine ten-eleven translocation dioxygenase (TET1) to specific promoters for the removal of the 5-methylcytosine (5mC) epigenetic mark, thereby modulating the epigenetic state of ESCs (54). Our proteomic data confirmed TET1 as an ESC-specific partner of LIN28A, further supporting the above-reported function. Among other epigenetic regulators, we identified core and accessory subunits of the Polycomb repressive complex 2 (PRC2) as LIN28A interactors in undifferentiated ESCs. PRC2 deposits H3K27me3 on histone H3, an epigenetic modification essential for fine-tuning transcriptional programs during development (51,55,56). PRC2, along with PRC1, plays a central role in silencing lineage-specific genes in pluripotent cells, repressing pluripotency genes and regulating key developmental genes as cells commit to specific lineages (50,51,57).

In ESCs and pre-implantation embryos, PRC2 is enriched at bivalent genes, which are chromatin domains marked by both H3K4me3 and H3K27me3 (58). These domains include lineage-specific genes, whose premature activation is prevented by PRC2 in early development and undifferentiated ESCs (59). Upon differentiation, this bivalent state must be resolved towards activation or repression to support lineage-specific transcriptional programs (46). Activation requires the eviction of PRC2 from nucleosomes, selectively removing H3K27me3 from lineage-specific genes (60). Numerous studies have supported the idea that this selective eviction is mediated *in cis* by nascent mRNA, that regulates its own production by removing PRC2 from the promoter, thereby further promoting gene transcription, and *in trans* by the interactions with some RBPs and ncRNAs (25–28,52,61–65). Although the direct binding of PRC2 to RNA remains debated (62,66–70), evidence suggests that PRC2 activity may rely indirectly on RNA via protein-protein interactions with RBPs (26,72,73).

Our findings show that, in neuronal differentiation, LIN28A absence impairs the proper distribution of PRC2 on the chromatin accompanied by increased H3K27me3 levels despite unchanged EZH2 protein levels. We propose that LIN28A modulates PRC2-chromatin association and its eviction to regulate epigenetic programs during neural differentiation. In the absence of LIN28A, PRC2 remains bound to chromatin, repressing neural differentiation genes while maintaining pluripotency gene expression (**Figure 6E**). We have shown that in Lin28a KO differentiating cells EZH2 occupancy is enriched, widespread and persistent on promoters of genes involved in neuronal differentiation (**Figure 5**) and that EZH2 enzymatic inhibition does not rescue the differentiation phenotype of Lin28a KO (**Figure 4E**). These results combined with the observations that the PRC2 binding on chromatin stimulates K27me3 deposition (48,74,75) and that EZH2 enzymatic inhibition does not efficiently displace PRC2 from the chromatin (33), support our idea that LIN28A influences PRC2 binding to chromatin rather than the enzymatic activity of the complex.

Our data support the eviction model of PRC2 regulation, potentially reconciling various perspectives on PRC2/RNA interactions. Indeed, we have found that LIN28A interaction with PRC2 is RNA-dependent, with the ncRNA Neat1 at least partially responsible for mediating this interaction. A recent study shows that during ESC-to-EpiLC transition, Neat1 absence disrupts PRC2 chromatin balance, impairing developmental gene repression and differentiation (26). LIN28A may utilize Neat1 as an RNA scaffold for the interaction with PRC2, facilitating PRC2 chromatin eviction and activation of differentiation genes. Further investigations are required to determine whether LIN28a influences nascent mRNA availability at PRC2 target genes.

Our findings suggest that LIN28A interaction with PRC2 has a selective role in ectodermal lineage differentiation. Indeed, Lin28a KO cells do not show evident defects in mesodermal and endodermal differentiation. Furthermore, Lin28a levels increase during neural differentiation while decreasing during mesendodermal lineage formation. Finally, ectopic LIN28A expression during endodermal differentiation does not alter PRC2 binding to chromatin (data not shown). These observations align with the emerging evidence that PRC2 recruitment varies by developmental stage, and it is guided by interactions with accessory proteins that are dynamically expressed during development in a cell-specific fashion (51). We propose that LIN28A contributes to conferring specificity to PRC2 in neuronal differentiation while leaving open the possibility that the Lin28a KO phenotype may reflect combined effects on epigenetic and translational control, as suggested by the cytoplasmic localization of LIN28A in ESCs.

In conclusion our findings establish a new RNA-dependent functional connection between LIN28A and PRC2 in regulating epigenetic state during ESC differentiation.

## Data availability

Raw data from proteomic analysis have been deposited in the ProteomeXchange Consortium with a dataset identifier PXD049332. ChIP-seq data have been deposited in the NCBI GEO database with the accession number GSE282208.

## Acknowledgements

The authors would like to thank: Tiziana Parisi and Tommaso Russo for the critical reading of the manuscript, the Microscopy Facility of DMMBM (Naples, Italy); Ludovica D’Auria and Caterina Missero of the Advanced Light Microscopy Facility of CEINGE (Naples, Italy) for help with imaging; Fondazione IRCCS Ca’ Granda Ospedale Maggiore Policlinico (Vittoria Moretti) and Istituto Nazionale Genetica Molecolare (Marco Ghilotti) for the sequencing.

## Author contributions

SPi performed experimental design, data collection, analysis and interpretation and contributed to manuscript writing and editing. EC, GD, LDL, VR, MZ, MM, EPS performed data collection, analysis and interpretation and contributed to manuscript editing. CD, GL, DSM, CL, PM, AS and FP performed data analysis and interpretation and contributed to manuscript writing and editing. SPa performed conceptualization and experimental design, data collection, analysis and interpretation, resources, project administration, writing.

---

**Supplementary Figure 1: Generation of Lin28a KO cells.**

A) Schematic representation of the sgRNAs used to generate Lin28a KO cells by CRISPR-CAS9 technology.

B) Immunofluorescence analysis of WT and Lin28a KO ESCs to verify the absence of LIN28A expression in two clones obtained using the 2 sgRNAs. Scale bars 50 μm. DAPI was used to counterstain the nuclei.

C) Western blot and immunofluorescence analysis of WT and Lin28a KO ESCs differentiated into SFEBs to verify the absence of LIN28A expression in two clones obtained using the two sgRNAs, upon 4 days of differentiation. GAPDH was used as loading control. Scale bars: 50 μm. DAPI was used to counterstain the nuclei.

**Supplementary Figure 2: Lin28a KO does not impair the maintenance of ESC undifferentiated state.**

A) Western blot analysis of endogenous LIN28A in WT and Lin28a KO ESC clones obtained with 2 different sgRNAs (#1 and #2). OCT4 was used as a pluripotency marker. GAPDH was used as loading control.

B) Alkaline phosphatase staining of WT and Lin28a KO ESCs. Both cell types were plated clonally and after 7 days in ESC medium were stained for AP activity. The number of AP-positive colonies of three independent biological replicates is reported in the graph, p value not significant.

C) Immunofluorescence analysis of pluripotency markers in WT and Lin28a KO undifferentiated ESCs. DAPI was used to counterstain the nuclei. Scale bars: 50 μm. The graph represents the number of positive cells counting >400/cells capturing 30 fields from three different passages of cells.

D) Markers of undifferentiated ESCs (Oct4, Nanog, Essrb) were measured by qPCR (n≥3, p value not significant).

E) Western blot analysis of pluripotency marker expression in WT and Lin28a KO ESC clones and relative quantitation (n=4, p value not significant). GAPDH was used as loading control.

**Supplementary Figure 3: Analysis of Lin28a KO cells.**

A) WT and Lin28a KO cells were transfected with the vector encoding a Flag-tagged form of LIN28A or the empty vector (Mock) and the differentiation into neural precursors was evaluated by qPCR analysis. n=4, *p≤0.05, ** p≤0.005.

B) WT and Lin28a KO ESCs were induced to differentiate into mesodermal lineages and pluripotency (Oct4) and differentiation (Brachyury,T) markers were analyzed by qPCR at different time points (day 0 and day 5) of differentiation. The differentiation into cardiomyocytes was evaluated by counting the EBs that showed beating hearts at 12 days of differentiation. n=4, *p≤0.05, ** p≤0.005.

C) WT and Lin28a KO ESCs were induced to differentiate into endodermal lineages and pluripotency (Oct4) and differentiation (Gata4 and Foxa2) markers were analyzed before and after (8 days) of differentiation by qPCR to evaluate proper differentiation. n=4, *p≤0.05, ** p≤0.005.

**Supplementary Figure 4: Functional and association analysis of LIN28A-interacting proteins identified in ESCs and SFEBs.**

A) Metascape-based enrichment analysis for biological processes of the ESC- or SFEB-specific LIN28A protein interactors identified in this study.

B) FunRich-based enrichment analysis for molecular function of the ESC-or SFEB-specific LIN28A protein interactors identified in this study.

C) STRING analysis of the mitochondrial gene expression proteins identified as common LIN28A-interacting species in ESCs and SFEBs. Only high-confidence interactions (interaction score 0.7) are shown.

D) STRING analysis of the ESC-specific LIN28A-interacting proteins identified in this study. Only high-confidence interactions (interaction score 0.7) are shown.

E) STRING analysis of the proteins related to chromatin remodeling from the LIN28A-interacting components uniquely identified in ESCs (medium-confidence, strength 0.53, False discovery rate 1.30 e-10).

**Supplementary Figure 5: EZH2 levels in Lin28a KO cells.**

A) Western blot analysis to compare the level of EZH2 in WT and Lin28a KO ESCs. The graph represents the mean ± SD of six independent experiments.

B) Reat time analysis of Ezh2 mRNA levels in WT and Lin28a KO cells. n=6. Ns: t-test not significant.

**Supplementary Table S1. Reagents and tools table.**

List of reagents and tools used throughout this study.

**Supplementary Table S2. List of LIN28A interactors identified by Mass Spectrometry.**

Protein identification results derived from mass spectrometry data, processed using Proteome Discoverer software. The following columns are shown: samples, protein false discovery rate (FDR) confidence, gene name, protein description, exp. q-value, Sum posterior error probability (PEP) score, sequence coverage (%), Mascot identification score values.

**Supplementary Table S3. Comparison of identified LIN28A interactors with CrapOME database and previous studies.**

LIN28A interactors identified in this study compared with CRAPome database or described in recent studies, and BioGRID, IntAct and STRING LIN28A interactome databases. Sample, gene name, protein description, number and percentage of entries matched in CRAPome database, found (X) in the paper (11,21), BioGRID (36), IntAct (37), and STRING database, number of finding in previous studies are reported.

**Supplementary Table S4. Genes mapped on the peaks identified by ChIP-seq in WT and Lin28a KO ESCs and SFEBs.**

List of EZH2 target genes mapped on the peaks identified in the ChIP-seq for EZH2 with a TSS within +/- 1000 bp of distance.

