## Supplementary figures and images for "The RNA binding protein LIN28A mediates chromatin dynamics during neuronal differentiation"

### Supplemental Figure 1

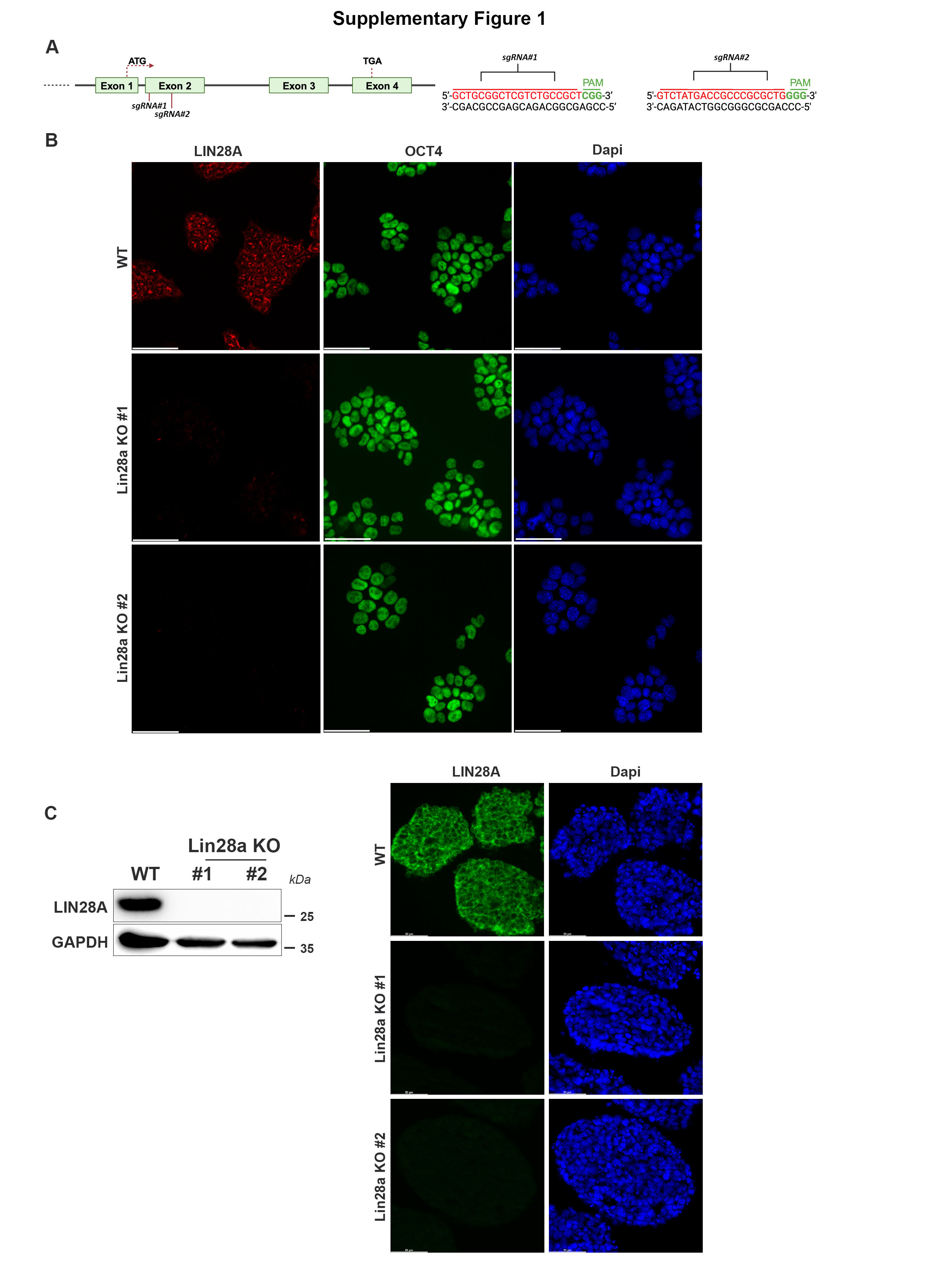

### Supplemental Figure 2

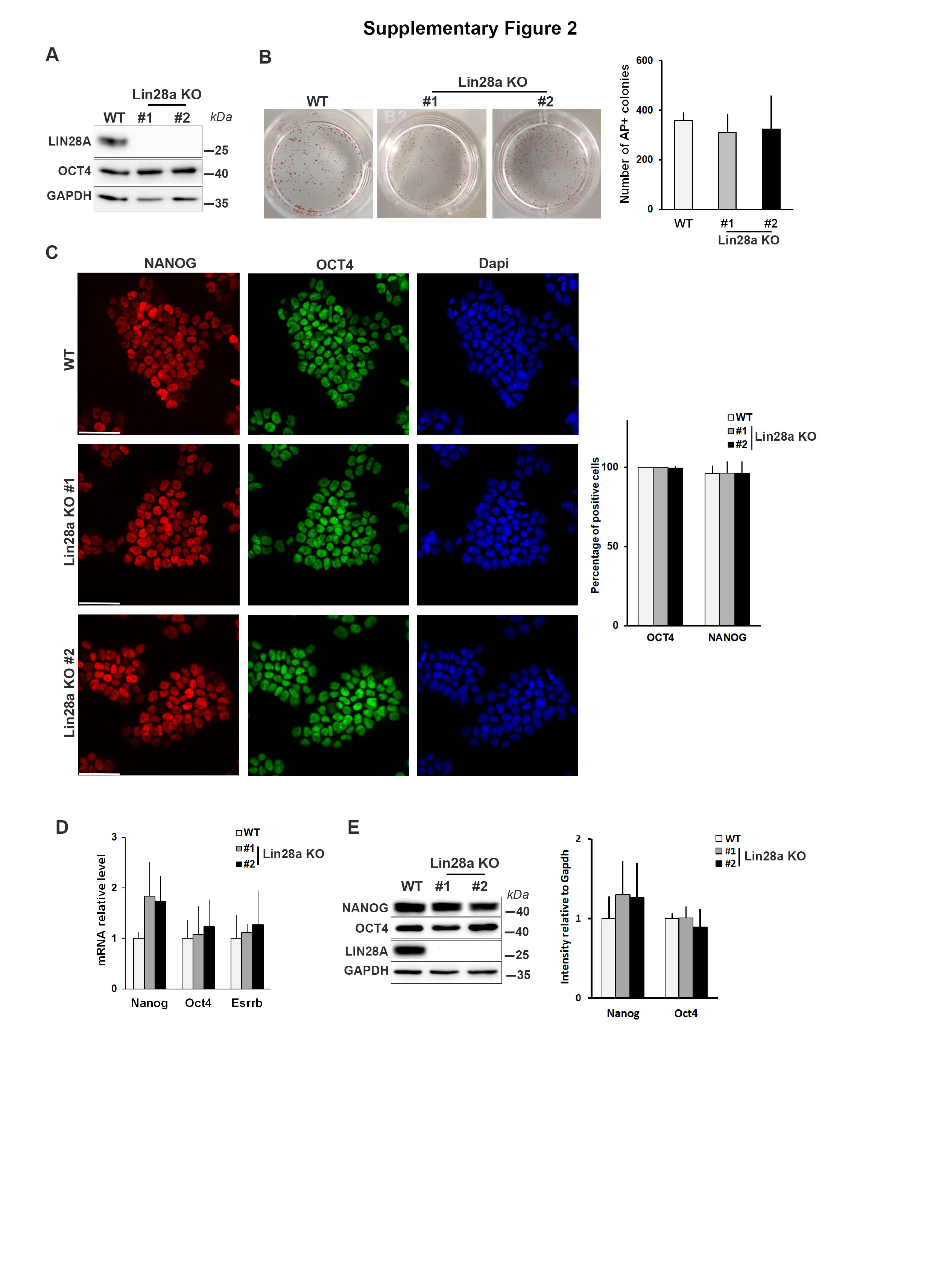

### Supplemental Figure 3

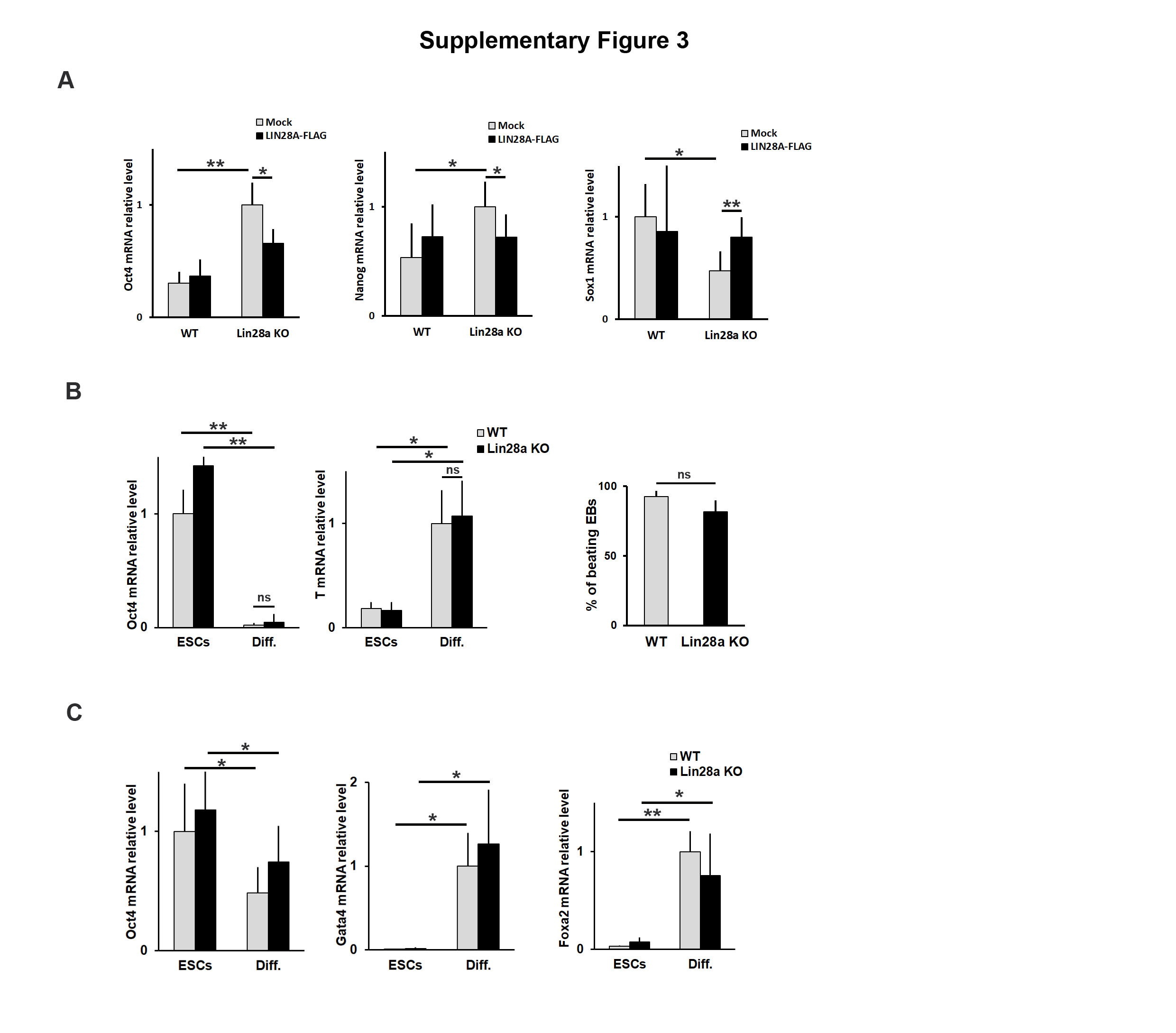

### Supplemental Figure 4

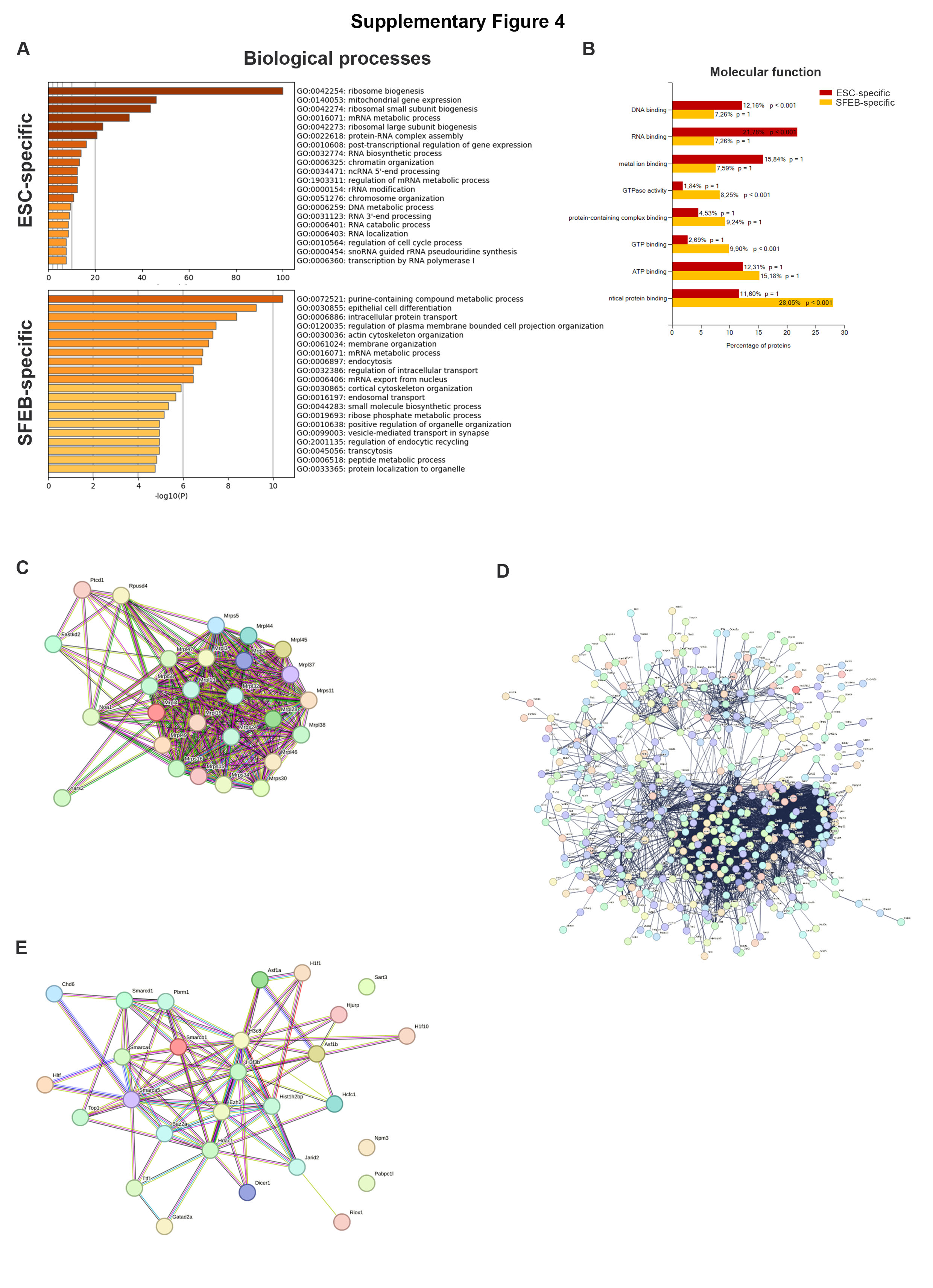

### Supplemental Figure 5

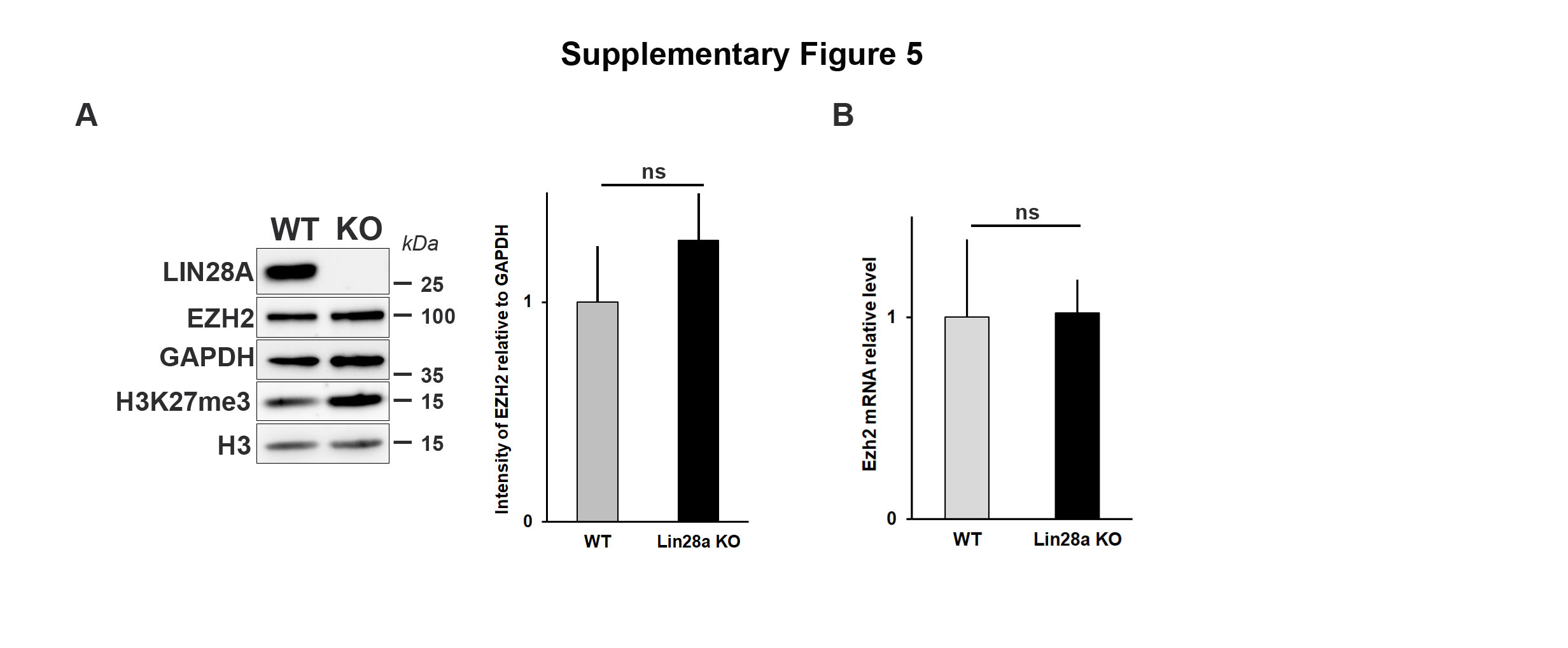
